# Defining the high-translational readthrough stop codon context

**DOI:** 10.1101/2024.12.20.629686

**Authors:** Daniela Smoljanow, Dennis Lebeda, Julia Hofhuis, Sven Thoms

## Abstract

Translational termination is not entirely efficient and competes with elongation, which might result in translational readthrough (TR). TR occurs when a near-cognate tRNA binds to a stop codon, (mis)interpreting it as a sense codon and producing a C-terminal extension of the protein. This process is influenced by the stop codon itself and the surrounding nucleotide sequence, known as the stop codon context (SCC). To investigate the role of these cis-acting elements beyond the high-TR motif UGA CUA G, this study examines specific positions within the SCC, both upstream and downstream of the motif, that contribute to variations in basal and aminoglycoside-induced TR. In particular, we identified a surprisingly large influence of the upstream nucleotide positions -9 and -8 (relative to the stop codon) and positions +11 and +12 on readthrough levels, revealing a complex interplay between nucleotides in the expanded SCC. These findings support our understanding of translational termination and may benefit the development of pharmacological therapy for diseases caused by premature stop codon mutations.

## Introduction

Translation of mRNA is catalyzed and coordinated by ribosomes and usually terminates when a stop codon enters the ribosomal A site. Eukaryotic cells universally recognize the three stop codons UAA, UAG, and UGA. Upon introduction of a stop codon in the ribosomal A site, the eukaryotic release factor 1 (eRF1) binds to the stop codon via its N-terminal domain [1, 2]. In eukaryotic cells, eRF1 recognizes all three stop codons, while the class II GTPase eRF3 facilitates termination through GTP hydrolysis, releasing the nascent polypeptide chain [3–5]. Termination is highly accurate, surpassing 99.9% efficiency [6, 7]. However, a near-cognate tRNA (nc-tRNA) with two complementary nucleotides can compete with eRF1 for stop codon binding, which may lead to decoding of the stop codon as a sense codon, a process known as translational readthrough (TR). This results in C-terminally extended proteins. This competition is influenced by the availability and properties of tRNAs, including their abundance, modifications, and affinity to specific codons [8–14].

The three stop codons exhibit different termination efficiencies, with UGA being the least efficient and having the highest TR rate, followed by UAG and UAA [6, 15]. The efficiency of nc-tRNA binding can be influenced by the nucleotide immediately following the UGA stop codon. The presence of a ’C’ right after UGA crucially impacts the nc-tRNA binding [11, 16, 17]. This demonstrates, that the fourth base is important for regulating stop codon recognition and translation termination efficiency [18, 19]. Crosslinking experiments have shown the interaction of eRF1 with the +4 nucleotide immediately following the stop codon [34]. In addition, further nucleotides near the stop codon are known to influence the TR efficiency [16, 20–23]. Both the 5’ and 3’ nucleotides of stop codon contexts (SCC) contribute significantly to readthrough efficiency and can lead to varying levels of extended protein production [14, 24, 25]. Data from machine learning analysis of readthrough efficiency from HEK293T ribosome profiling experiments, show evidence for the conservation of the stop codon context and 3’-UTR length in regulating TR [14].

TR can exhibit functional activity within the cell if the C-terminally extended protein acquires a feature that differs from that of the parental protein, a mechanism known as functional TR (FTR) [26–28]. FTR increases the proteome’s functional capacity without extending the genome. The intricate interplay between precise termination and TR underscores the complexity of translational regulation, allowing the synthesis of diverse protein isoforms from the same mRNA template [10, 17, 26, 27].

FTR was first described for viral genes [20, 29, 30]. Phylogenetic analysis and ribosome profiling in Drosophila and other metazoa revealed a large number of genes involved in TR [31, 12, 32]. FTR has also been found in mammals for some genes [32, 11, 33]. Using an in silico regression model, our group identified a TR-promoting nucleotide context in 57 human genes, with experimental validation confirming TR in six of them and measuring that in different cell types. Precisely, we identified the nucleotide consensus sequence CUA G downstream of the UGA stop codon for high TR propensity [17], which we call the high-TR motif. Deletion experiments showed the CUA G as a key determinant of TR efficiency [11].

Notably, positions outside the high-TR contexts impact TR efficiency. Specific sequences downstream of the stop codon, including the +7, +8, and +9 positions, also determine TR efficiency in yeasts, potentially by forming secondary structures [22]. Furthermore, the immediate nucleotides upstream of the stop codon have been shown to modulate TR in bacteria and yeasts [34] and also in mammalian cells. In experiments conducted on mouse fibroblasts (NIH3T3) and HEK293T cell lines, the highest TR levels were observed when the first position upstream of the termination codon was occupied by adenine or guanine, while uracil at this position is associated with lower TR levels [11, 35]. In yeast, adenine at positions −1 and −2 enhances TR, particularly for the UAG stop codon, but also for other termination codons [36, 37].

The importance of the nucleotide composition surrounding the stop codon, both 3’ and 5’, is underscored by their impact on TR efficiency in various FTR transcripts, such as *Malate dehydrogenase 1* (*MDH1*), *Aquaporin 4 (AQP4)*, and *Lactate dehydrogenase B* (*LDHB*) affecting the C-terminal extension of the expressed proteins [17, 28, 33]. Due to the presence of the high-TR motif, all these transcripts undergo FTR but exhibit markedly different TR efficiencies. Both TR-extended isoforms MDH1x and LDHBx (x for extended) gain a peroxisomal targeting signal (PTS1) for peroxisomal translocation [17, 33, 28]. MDH1 is essential for redox balance, in the citric acid cycle and the malate-aspartate shuttle. In the peroxisome, MDH1x and LDHBx assist in the regeneration of redox equivalents. A recent study showed that MDH1x regulates NAD(H) levels in peroxisomes [38]. The readthrough-extended Aquaporin 4 variant (AQP4x) is specifically localized at astrocytic endfeet surrounding brain blood vessels and modulates the supramolecular organization of AQP4 in astrocytic membranes [39, 40].

A comprehensive understanding of the SCC and its impact on TR efficiency is crucial for advancing our knowledge of translational control. This study investigates SCC’s influence by analyzing nucleotide contributions at key positions, focusing on FTR transcripts *MDH1*, *AQP4*, and *LDHB.* Despite the same high-TR motif, the reason for their different TR levels remains unclear. By interchanging upstream and downstream sequences beyond the high-TR motif and systematically exchanging specific nucleotide positions between *MDH1* and *LDHB* we detect a so far underestimated influence of the upstream and downstream sequence beyond the high-TR motif.

## Results

### The influence of the 5’ context on translational readthrough efficiency surpasses that of the 3’ context beyond the CUA G motif

*MDH1*, *AQP4,* and *LDHB* share the same high-TR motif but exhibit different levels of readthrough efficiency. To evaluate the TR efficiency, we used a dual reporter assay, ensuring independence from variations in mRNA level expression as previously established [41]. This reporter system encodes an N-terminal tagRFP and a C-terminal eGFP, separated by the SCC which encompasses nucleotide positions -10 to +13 including the stop codon at position +1 to +3 (Figure 1A). The system is based on a high-content flow cytometry measurement of transiently transfected HeLa cell suspensions in a 96-well plate [42]. TR was determined by evaluating the ratio of eGFP to tagRFP fluorescence, with normalization against a 100% TR control, a construct without a stop codon in the SCC for continuous translation of eGFP. This construct serves as a reference point for maximal eGFP fluorescence.

**Figure 1:**
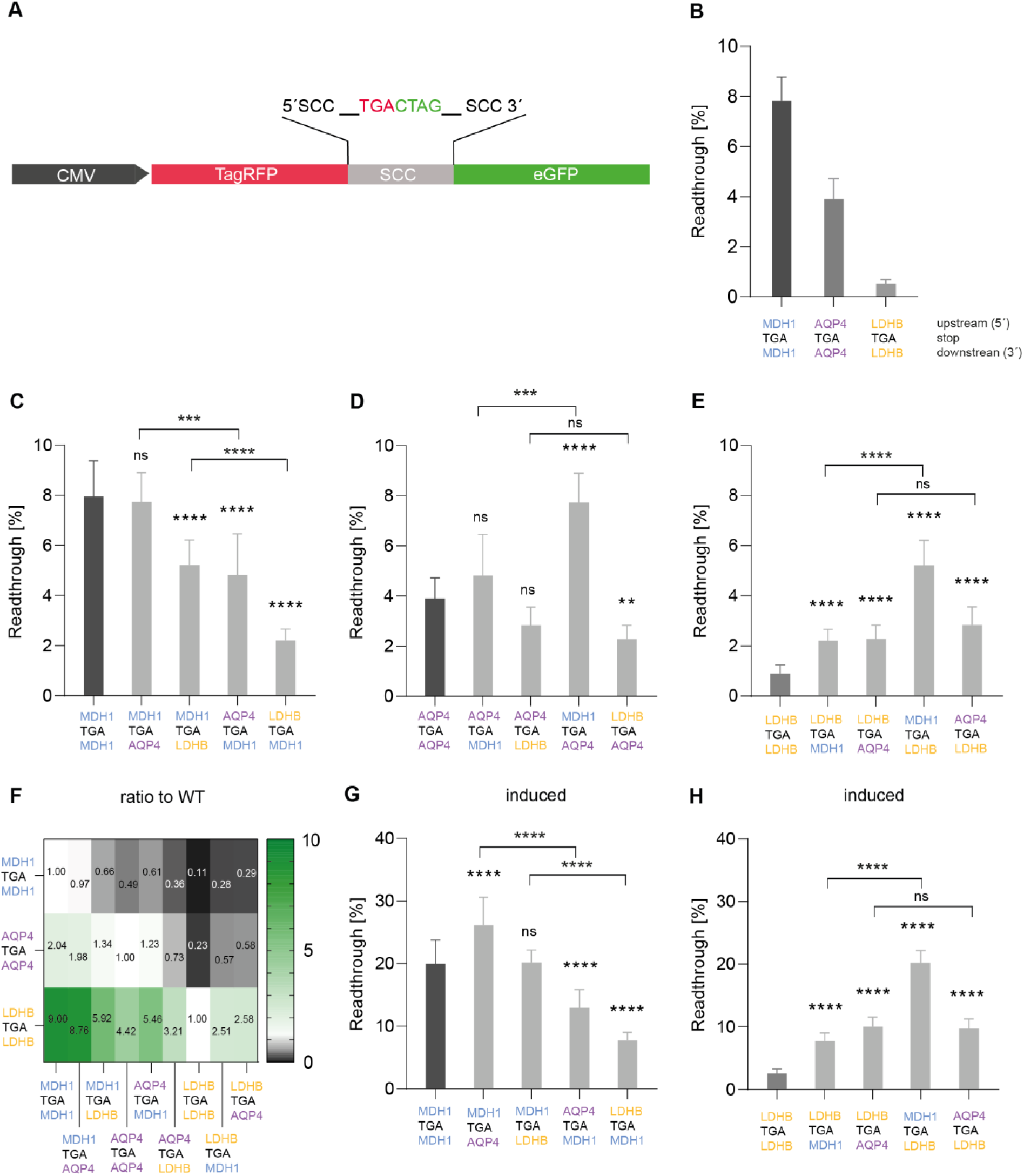
The stop codon context influences both basal and induced translational readthrough. The upstream sequence has the next largest impact beyond the high TR motif. **A)** Schematic Representation: The SCC is located between a 5’ tagRFP and a 3’ eGFP. If TR occurs, a protein with both RFP and GFP is formed. TR is calculated as GFP over RFP fluorescence. **B)** Basal TR of SCCs: TR of SCCs from WT *MDH1*, *AQP4*, and *LDHB* in HeLa cells, was measured in three experiments with three replicates (n=9). TR was measured using a dual reporter assay (5’ RFP, SCC, 3’ GFP). Plasmid PST1596 (no SCC) served as a 100% TR control. **C-E)** Basal TR Efficiency: TR efficiency of WT SCC sequences of *MDH1*, *AQP4*, and *LDHB*, and cross-combined SCC constructs. Measurements were done in HeLa cells with at least three experiments and three replicates. Data normalized to PST1596 control. **C)** WT sequence of *MDH1* with hybrid SCC constructs (*MDH1/AQP4* and *MDH1/LDHB*). **D)** WT sequence of *AQP4* with hybrid SCC constructs (*AQP4/MDH1* and *AQP4/LDHB*). **E)** WT sequence of *LDHB* with hybrid SCC constructs (*LDHB/MDH1* and *LDHB/AQP4*). **F)** Heat Map: Ratios of basal TR levels of WT SCCs (*MDH1, AQP4, LDHB*) and cross-combined SCC constructs compared to WT reference. Values are presented as mean with standard error. Statistical analysis was performed using one-way ANOVA and Dunnett’s multiple comparison test. **G-H)** Induced TR of SCCs: Comparison of induced TR in WT *MDH1* and *LDHB* with hybrid SCC constructs (*MDH1, AQP4, LDHB*). Measured in HeLa cells with at least three experiments, three replicates each, with and without induction. Data normalized to PST1596 control.

We first tested the wild-type SCCs of *MDH1, AQP4,* and *LDHB* in HeLa cells to validate differences between these sequences. The results showed a TR efficiency of 7.8% for the *MDH1* SCC. In contrast, the *AQP4* SCC exhibited a lower TR efficiency of 3.9%, indicating a 2-fold difference between *MDH1* and *AQP4*. Notably, the *LDHB* SCC TR efficiency with 0.5% showed the lowest TR level, which is a > 15-fold difference compared to *MDH1*, and a > 7-fold difference compared to *AQP4*. Despite all transcripts harboring the same stop codon and the CUA G motif known to promote TR, they show varying TR levels (Figure 1B).

To investigate the relative impact of the upstream (-10 to -1) and downstream SCC (+4 to +13), we generated cross-combined constructs with the upstream sequence of one transcript combined with the downstream sequence from another transcript, all derived from *MDH1*, *AQP4*, and *LDHB.* Compared to the wild-type *MDH1* SCC with 7.8%, the cross-combined SCC with the upstream *MDH1* and the downstream *AQP4* sequence showed no significant change in TR efficiency, whereas the reverse arrangement led to a significant decrease in the basal TR level to 4.8% (Figure 1C), which corresponds to approximately 60% of the wild-type (Figure 1F). Similarly, combining the upstream *MDH1* sequence with the downstream *LDHB* sequence resulted in a significant decrease in TR efficiency to 5.2% (∼0.7-fold wild-type *MDH1*). Remarkably, the opposite arrangement of the upstream *LDHB* sequence and downstream *MDH1* sequence led to a further significant decrease in TR efficiency to 2.2% (Figure 1C) and a ratio of ∼0.3 times that of wild-type (Figure 1F).

The basal TR efficiency of wild-type *AQP4* increased from 3.9% to 7.7% after the replacement of the upstream SCC with *MDH1* (Figure 1D). Compared to the wild-type, the results indicate a ∼2-fold increase (Figure 1F). On the other hand, the opposite arrangement increased efficiency by 4.8%. However, combining the upstream *LDHB* sequence with *AQP4*’s downstream sequence decreased TR efficiency to 2.3% (Figure 1D). Exchanging the downstream SCC of *AQP4* with *LDHB* did not alter basal TR efficiency compared to *AQP4* wild-type.

For constructs containing parts of *LDHB* sequence compared to wild-type *LDHB* SCC, all replacements led to increased TR levels. Notably, the basal TR efficiency of *LDHB* SCC enlarged from 0.9% to 5.2% (Figure 1E), a ∼5.9-fold increase when the upstream *MDH1* sequence was combined with the downstream *LDHB* (Figure 1F). The replacement of the downstream sequence from *LDHB* with *MDH1* showed an increase to 2.2%, a ∼2.5-fold change (Figure 1E, F). Fusion of the upstream *AQP4* sequence with the downstream *LDHB* sequence increased efficiency to 2.8%, while the reverse combination increased the TR efficiency to 2.3% (Figure 1E).

In summary, the ‘best’ SCC (*MDH1*) cannot be improved, while the ‘mid’ SCC (*AQP4*) can be improved only by the 5’ *MDH1* SCC, whereas the ‘weakly’ performing (high-TR) SCC *LDHB* is improved by any exchange, but received the highest increase by the 5’ *MDH1* SCC. Consistent with recent findings, we observed a greater influence of the 5’ sequence on TR efficiency than the 3’ sequence [43] (Figure 1C-F).

Aminoglycosides like geneticin (G418) are known to induce TR by promoting the misreading of stop codons as sense codons [44, 45]. Our findings demonstrate that SCCs exhibiting high basal TR levels also respond effectively to induction by G418. Induction of wild-type *MDH1* SCC led to a TR efficiency of ∼20%, a 2.6-fold increase over the basal level. G418 raised the TR efficiency to 26.2% when the downstream *MDH1* sequence was exchanged by the respective *AQP4* sequence. In contrast, the upstream *MDH1* sequence combined with the downstream *LDHB* sequence showed no significant effect (Figure 1G). Both the upstream *AQP4* sequence and the upstream *LDHB* sequence combined with the downstream *MDH1* sequence resulted in a reduction of the TR level (Figure 1G). For *LDHB* wild-type SCC, G418 induced the TR to 2.5% (Figure 1H). Overall, a value of 20% TR efficiency was measured for the combination of the upstream *MDH1* sequence and the downstream *LDHB* sequence with G418-induction (Figure 1H), which exhibited an 8-fold increase in TR efficiency compared to the wild-type. The fusion of the upstream *AQP4* sequence with the downstream *LDHB* sequence led to a TR efficiency level of 9.8% (Figure 1H).

Altering the upstream sequence from *MDH1* to *AQP4* or *LDHB* in cross-combined constructs consistently has a large impact on TR efficiency for both basal and inducible TR, highlighting the upstream SCC’s dominant role.

### Decoding the distinctive impact: Disparate effects of double and single nucleotide exchanges on translational readthrough efficiency

*MDH1* and *AQP4* showed higher basal TR efficiencies than *LDHB*. *MDH1* and *AQP4* share the high-TR motif (Figure 2A, highlighted in green) and further eight SCC nucleotides (Figure 2A, highlighted in blue/purple), that differ in the *LDHB* SCC (Figure 2A, highlighted in yellow). To analyze whether these nucleotides are responsible for the strong effect on TR efficiencies, we systematically exchanged individual nucleotides and nucleotide pairs between *MDH1* and *LDHB* (Figure 2A).

**Figure 2:**
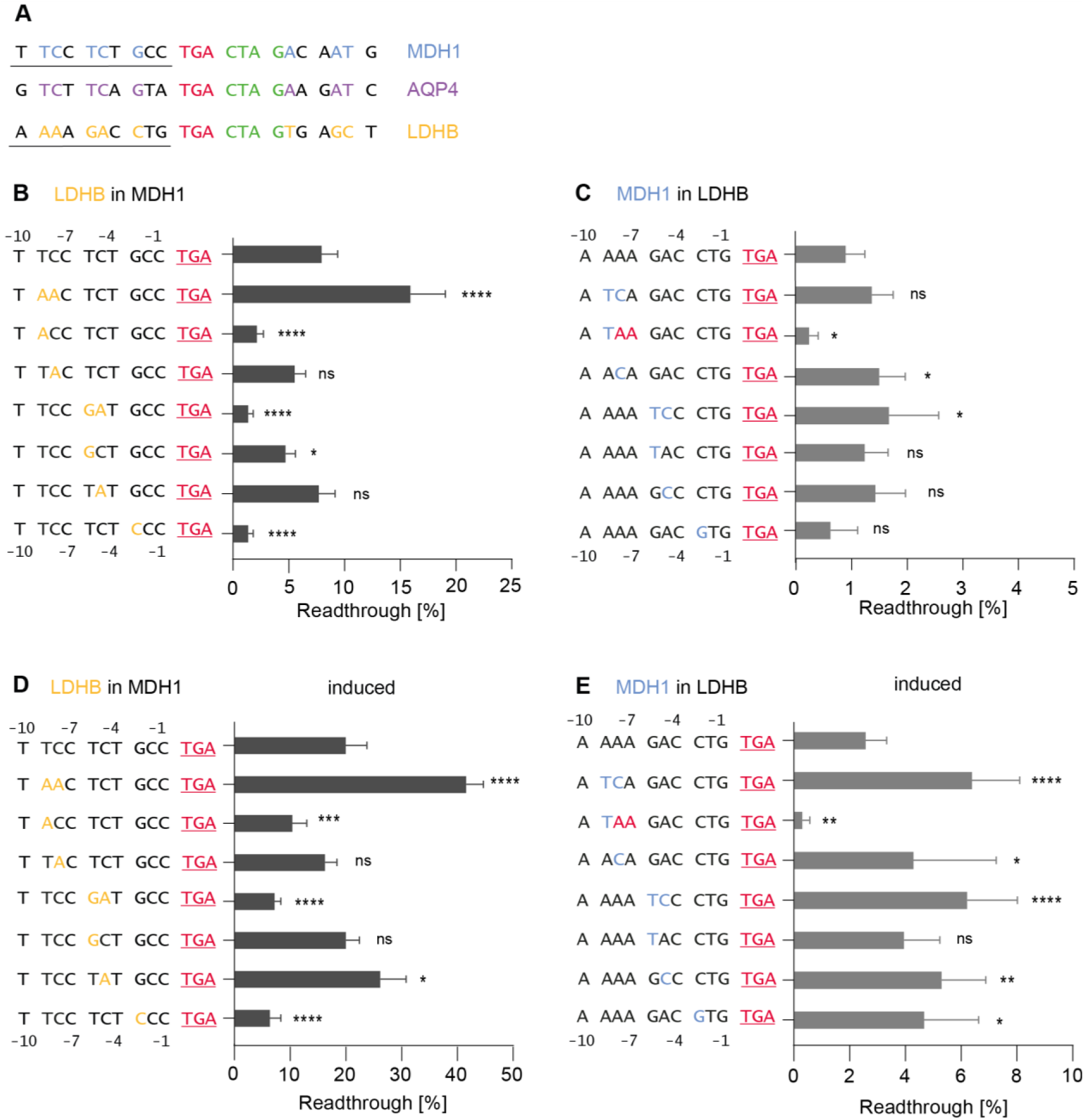
Both single and double nucleotide substitutions within the stop codon context lead to changes in translational readthrough efficiency for both basal and inducible readthrough. Single exchanges in the upstream sequence do not exhibit the same tendencies as double exchanges. **A)** SCC (-10 to -1_TGA_+4 to +13): *MDH1*, *AQP4*, and *LDHB* transcripts. The stop codon is in red. Green indicates nucleotides downstream of the stop codon promoting TR (high TR motif). Blue and purple show nucleotides shared between *MDH1* and *AQP4.* Yellow marks *LDHB*-specific nucleotides where *MDH1* and *AQP4* share the same nucleotides. **B-C)** Basal and induced TR of *MDH1* SCC with altered upstream positions: B) Basal TR of WT *MDH1* SCC and constructs with individual nucleotide exchanges. Nucleotides from *MDH1* were replaced with corresponding *LDHB* nucleotides. **C)** Induced TR with G418: Induced TR of WT *MDH1* SCC and constructs with individual nucleotide exchanges. **D-E)** Basal and induced TR of *LDHB* SCC with altered upstream positions: **D)** Basal TR of WT *LDHB* SCC and constructs with individual nucleotide exchanges. Nucleotides from *LDHB* were replaced with corresponding *MDH1* nucleotides. **E)** Induced TR with G418: Induced TR of WT *LDHB* SCC and constructs with individual nucleotide exchanges. Black letters show the wild-type *MDH1* and *LDHB* sequences. Yellow letters indicate nucleotides from *LDHB* SCC, and blue letters indicate nucleotides from *MDH1* SCC. Measurements were conducted at least three times with three replicates. Data normalized to a 100% TR control. Statistical analysis was performed using one-way ANOVA and Dunnett’s multiple comparison test.

We first investigated the upstream sequence. Surprisingly, an exchange of *MDH1* TC at positions -9 and -8 with *LDHB* AA led to a ∼2.2-fold increase in TR efficiency (Supplementary Figure 1A) from 7.8% to 16% (Figure 2B). In contrast, the individual exchange of position -9 resulted in a significant decrease in TR efficiency (2.2%), while the replacement at position -8 alone showed no effect. Consequently, the combined exchange at positions -9 and -8 and the single exchanges at position -9 exhibited distinct alterations in the TR efficiency in opposite directions (Figure 2B, Supplementary Figure 1A). The combined exchange in the *MDH1* SCC at positions -6 and -5 with *LDHB* nucleotides TC→GA decreased TR efficiency to 1.4% (Figure 2B). Similarly, a single exchange at position -6 with T→G yielded a TR efficiency of 4.7%. Replacing the -3 position also significantly decreased TR efficiency to 1.4% (Figure 2B).

Next, we replaced *LDHB* nucleotides within the SCC with *MDH1* nucleotides. The basal TR efficiency of the wild-type *LDHB* SCC was 0.9%. Surprisingly, the combined exchange of *LDHB* nucleotides with *MDH1* nucleotides at positions -9 and -8 (AA→TC) showed no significant change in TR efficiency (Figure 2C). In contrast, when we replaced individual nucleotides, the -9 A→T nucleotide exchange decreased the TR efficiency to 0.3% due to the introduction of an additional in-frame stop codon. The A→C-exchange at position - 8 increased TR efficiency to 1.5% (Figure 2D, Supplementary Figure 1B).

Similarly, GA→TC exchange at positions -6 and -5 in *LDHB* increased TR efficiency from 0.9% to 1.7%. The respective single exchanges at these positions did not alter TR efficiency significantly (Figure 2C). An exchange at position -3 showed no significant alteration in TR efficiency and exhibited a ratio of ∼0.9 (Figure 2C; Supplementary Figure 1B). Small changes in specific upstream nucleotides can significantly alter TR efficiency, demonstrating that even closely related sequences can have dramatically different effects on TR.

Upon G418-induction, TR efficiency significantly increased for both *MDH1* and *LDHB* sequences. Notably, the TC→AA exchange at positions -9 and -8 in *MDH1* increased TR efficiency to 41.6% (2.6-fold compared to non-induced, 2.6-fold compared to wild-type induced; Figure 2D, Supplementary Figure 1A). The single exchange at position -9 in *MDH1* resulted in a higher TR efficiency of 10.4% upon induction but below the induced TR efficiency of the wild-type (Figure 2D).

Surprisingly, for *LDHB* constructs, induction led to TR efficiencies exceeding 6%, a substantial increase from the basal 0.9% (Figure 2E). Specific exchanges demonstrated varying responses to induction. For example, the GA→TC exchange at positions -6 and -5 in *LDHB* resulted in a G418-induced TR efficiency of 6.2%, higher than individual exchanges. This combined exchange exhibited a high ratio to the wild-type, surpassing the G418-induced single exchanges at these positions with ∼2.3 compared to the wild-type (Figure 2E and Supplementary Figure 1B).

Interestingly, double exchanges of nucleotides, which are located next to each other, did not always have the same effect as the corresponding single exchanges within the SCC, as observed for the *MDH1* sequence at positions -9 and -8. It is particularly noteworthy that the varying TR efficiencies appear to result from combinations of nucleotides rather than from individual nucleotides. Specific nucleotide exchanges significantly influenced TR efficiency. Ratios were calculated for G418-induced constructs, revealing consistent behavior across most compared to non-induced counterparts (Supplementary Figure 1). Importantly, while basal TR efficiency varies with nucleotide sequences, G418 induction amplifies these effects in a manner consistent with the baseline trends. The nucleotide exchanges that impact TR efficiency at basal levels show similar directional changes under G418 treatment, though with increased magnitude.

### Nucleotides at positions +11 and +12 significantly affect translational readthrough efficiency

To further investigate the influence of downstream nucleotides on TR efficiency, we examined substitutions in the downstream SCC, focusing on *MDH1* and *LDHB*. In the *MDH1* SCC, the A→T substitution at position +8 significantly increased basal TR efficiency to 14% (Figure 3A). Further investigation into *MDH1*’s downstream SCC revealed a more substantial enhancement with the AT→GC exchange at positions +11 and +12. Despite their distance from the stop codon, this alteration boosted TR efficiency to 18.9%, underscoring the profound impact of more distant nucleotides on TR efficiency. Individual substitutions at either +11 (A→G) or +12 (T→C) positions in *MDH1* did not yield significant changes in TR efficiency (Figure 3A).

**Figure 3:**
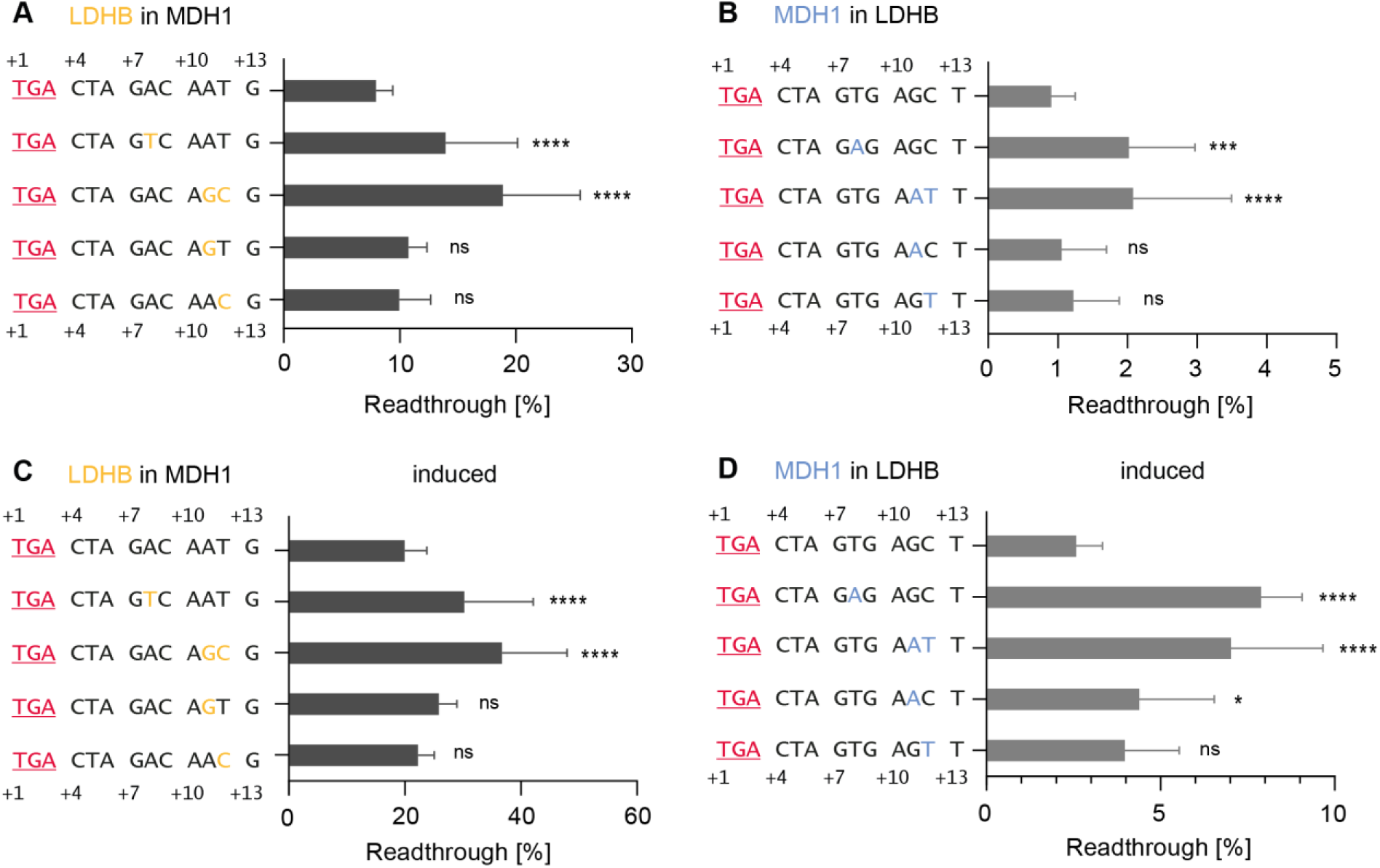
Both single and combined nucleotide exchanges, which are located next to each other, within the downstream stop codon context affect TR efficiency for both basal and inducible readthrough. Single exchanges in the upstream sequence do not display the same patterns as double exchanges. **A-B)** Basal and induced TR of *MDH1* SCC with altered downstream positions: **A)** Basal TR of WT *MDH1* SCC and constructs with individual nucleotide exchanges. Nucleotides from *MDH1* were replaced with corresponding *LDHB* nucleotides. **B)** Induced TR with G418: Induced TR of WT *MDH1* SCC and constructs with individual nucleotide exchanges. **C-D)** Basal and induced TR of *LDHB* SCC with altered downstream positions: **C)** Basal TR of WT *LDHB* SCC and constructs with individual nucleotide exchanges. Nucleotides from *LDHB* were replaced with corresponding *MDH1* nucleotides. **D)** Induced TR with G418: Induced TR of WT *LDHB* SCC and constructs with individual nucleotide exchanges. Black letters show the wild-type *MDH1* and *LDHB* sequences. Yellow letters indicate nucleotides from *LDHB* SCC, and blue letters indicate nucleotides from *MDH1* SCC. Measurements were conducted at least three times with three replicates. Data normalized to a 100% TR control. Statistical analysis was performed using one-way ANOVA and Dunnett’s multiple comparison test.

In the *LDHB* SCC, the T→A exchange at position +8 doubled the TR level to 2% (Figure 3B). Similarly, the double exchange at positions +11 and +12 with GC→AT exhibited an increase to 2.1% in TR efficiency, mirroring the effect seen with the +8-position exchange. Individual substitutions at positions +11 (G→A) and +12 (C→T) in the *LDHB* context showed no significant effect on TR efficiency (Figure 3B).

Following induction with G418, various constructs showed significant increases in TR efficiency. Specifically, the exchange at position +8 in *MDH1* with *LDHB* nucleotide led to a TR efficiency of 30.3% (Figure 3C). The double exchange at positions +11 and +12 (AT→GC) resulted in a substantial TR efficiency increase to 36.8%, significantly higher than induced wild-type *MDH1* SCC. In contrast, individual substitutions at positions +11 and +12 showed no significant alterations compared to wild-type *MDH1* SCC (Figure 3C). When calculating mutant versus wild-type TR ratios, values for basal TR ranged approximately between ∼1.2 and ∼2.7, whereas the induced ratios ranged between ∼1.0 and ∼2.3 (Supplementary Figure 1C).

Examining *LDHB* SCC and corresponding constructs, it was observed that only one construct did not significantly differ from the induced wild-type. The TR value of the wild-type *LDHB* sequence increased from basal 0.9% to 2.6% after induction with G418 (Figure 3D). Induction with G418 and the insertion of adenine instead of thymine at position +8 led to an increase in TR to 7.9%. The double exchange at positions +11 and +12 (GC→AT) resulted in a TR efficiency of 7%, and the single exchange at position +11 (G→A) led to a measurable G418-induced TR level of 4.4%. Similarly, the construct with the substitution at position +12 and the T→C-exchange yielded a G418-induced TR of 4% (Figure 3D). Mutant versus wild-type TR ratios indicate that the basal TR ratio for the +8 exchange increased nearly threefold compared to the reference (∼2.8). Similarly, the double exchange at positions +11 and +12 exhibited a ∼2.9-fold increase in TR efficiency compared to the wild-type (Supplementary Figure 1D).

Exchanges at positions +8, +11, and +12 reveal that distant nucleotides significantly influence TR efficiency, with double exchanges having pronounced effects. Systematic SCC modifications provide valuable data for in silico TR calculations, enhancing the predictability of FTR candidates and their TR efficiencies. Improved prediction tools can refine the search for FTR candidates.

## Discussion

In this study, we show the critical impact of distinct positions within the SCC on TR efficiency, highlighting the intricate interplay of nucleotide combinations within the sequence. Consequently, TR emerges as a finely regulated mechanism, prompting a thorough examination of the role of the 5’ SCC and the influence exerted by nucleotides positioned distally from the stop codon. Furthermore, the results emphasize the potential combinatorial effects of nucleotides on TR levels.

### The upstream stop codon context has the next largest impact after the high TR motif

To better understand the impact of the 5’ and 3’ SCCs on TR, wild-type SCCs of *MDH1*, *AQP4*, and *LDHB* were tested in HeLa cells, and the effects of the 5’ and 3’ SCCs were investigated by flow cytometry [42]. Combined constructs from *MDH1*, *AQP4*, or *LDHB* wild-type demonstrated that the 5’ SCC exerts a stronger influence on the TR efficiency in these SCCs than the 3’ SCC. We showed significant differences in both basal and induced TR efficiencies across 5’ SCC replacements compared to the corresponding wild-type SCC and some 3’ SCC exchanges.

Our results align with recent findings that the 5’ sequence has a substantial impact. The TR levels of *OPRL1* and *DUS4L* including 18 nucleotides upstream and twelve nucleotides downstream of the stop codon and the high-TR motif CUA G have been investigated by systematically swapping the upstream sequences. It was shown that the 5’ context accounts for a fourfold difference in TR efficiency compared to a twofold difference of the 3’ sequence. Specifically, the identity of the P-site codon directly preceding the stop codon was identified as a critical determinant of TR efficiency, highlighting the importance of sequence composition in shaping termination and readthrough outcomes [17, 43].

### Influence of individual nucleotides in the 5’ stop codon context and the combinatorial effect of nucleotides

So far, only a few studies in eukaryotes have analyzed the specific role of 5’ signals [34–36, 46]. Interestingly, the insertion of a cytosine at the *MDH1* position -3 led to a significantly lower TR efficiency, which aligns with findings in a case report with a patient with junctional epidermolysis bullosa, where a cytosine at position -3 acts as a determinant of TR efficiency [47].

Strikingly, replacements at positions -6 and -5 of both *MDH1* (TC→GA) and *LDHB* (GA→TC) showed a substantial effect on TR efficiency. The TR level of *MDH1* significantly decreased, whereas that of *LDHB* increased significantly, highlighting the importance of specific nucleotide positions in TR regulation, which aligns with an in silico regression model, that shows a positive contribution to TR efficiency [17] This suggests that TR efficiency is fine-tuned by the specific nucleotide composition at the SCCs.

Previous phylogenetic analysis showed that positions upstream of the stop codon exhibit an evolutionary sequence conservation in TR candidates. Three of five analyzed TR candidates showed conservation of adenines (AA) at the -9 and -8 positions in the genes *OPRK1, OPRL1,* and *SACM1L* in a phylogenetic analysis of 29 mammalian species [11]. Consistent with this, our findings show the presence of adenines (AA) at -9 and -8 in wild-type *LDHB*. Notably, introducing this AA motif into the upstream sequence of *MDH1* significantly increases the TR efficiency. This highlights a synergistic relationship between conserved upstream adenines and downstream TR-promoting sequences, such as CUA G. Alongside these findings, a single substitution of adenine at position -9 leads to a decrease in TR efficiency. Evolution does not maximize TR but rather achieves an optimal level that balances functional necessity with regulatory constraints.

Our study extends previous findings that emphasize the codon in the P site (-3 to -1) as a significant factor influencing TR efficiency. This indicates that variations in the 5′-sequence can be critical determinants of translation termination efficiency [43]. With this study, we show that the high-TR motif can be redefined. Furthermore, we show that a combined exchange at position +11 and +12 (AT→GC) in the *MDH1* SCC and (GC→AT) in *LDHB* SCC increases the TR efficiency, while the single exchanges do not alter the efficiency.

### The amino acids in the nascent polypeptide chain possibly influence the translational readthrough by interacting with the ribosomal exit tunnel

The process of translational decoding identifies the correct tRNA for an A-site codon with error rates as low as 10⁻³ to 10⁻⁴. Despite this precision, ribosomes maintain a fast decoding pace, of 5–20 amino acids per second. This efficiency relies on rapid tRNA screening and engagement with the ribosome [48]. Mutations in the upstream sequence of the stop codon, altering the nascent peptide, significantly influence TR efficiency. The ribosomal exit tunnel acts as a selective gate, interacting with the nascent peptide chain to regulate translation. These interactions may influence TR by modulating ribosome kinetics and stop codon recognition [49]. In the SCC of *MDH1,* the mutation at position -3 converts the codon GCC to CCC. This change in the mRNA sequence is responsible for the incorporation of proline instead of alanine, which could lead to a translation slowdown. The kinetic properties of amino acids during peptide bond formation play a critical role in determining the efficiency of translation. For example, due to the unique cyclic structure of proline, its incorporation into polypeptides is significantly slower than other amino acids like phenylalanine or alanine. Proline’s slower incorporation during translation is primarily due to steric constraints introduced by its cyclic pyrrolidine ring. This structure reduces the flexibility of Pro-tRNA and hinders its accommodation into the ribosome’s A-site. Additionally, steric hindrance limits the accessibility of the amino group to the peptidyl transferase center, further delaying peptide bond formation. These effects make proline incorporation 15–23 times slower than alanine or phenylalanine [49, 50]. The translation of polycationic peptides, such as poly-lysine sequences highlights how peptide charge, mRNA sequence, and RNA modifications influence the ribosome’s translation speed [51]. Experimental evidence suggests that peptide characteristics and codon context can independently modulate ribosome kinetics, with codon usage and peptide interactions at the ribosomal exit tunnel playing distinct roles in shaping translation speed and ribosome behavior. These findings indicate that the properties of the nascent peptide and the mRNA sequence act as separate but interconnected factors in regulating translational processes [51]. This provides a basis for exploring how upstream mutations, which alter both mRNA sequence and nascent peptide properties, might synergistically affect TR efficiency, ribosome stalling, and termination fidelity in future studies.

### Finetuning of the high-readthrough motif influences both, basal and inducible translational readthrough efficiency

Translational readthrough-inducing drugs (TRIDs) are being researched for their potential to bypass premature termination codons and recover full-length proteins. Genetecin (G418) is a known gold standard for experimental TR induction as it provides the properties that allow the ribosome to keep translating but disturb its proofreading function [52–54].

Our study demonstrates that an increase in basal TR in chimeric constructs composed of two transcripts also leads to enhanced inducible readthrough. Furthermore, both combined and single nucleotide exchanges that elevate basal TR levels show improved inducibility of TR with G418 and vice versa, which is particularly relevant for TR of PTCs. These findings suggest that TR therapy is more likely to succeed in the presence of such motifs, that show characteristics of high-TR motifs. In summary, our study demonstrates that the 5’ and 3’ sequences surrounding the stop codon are crucial for TR efficiency. The complex interplay of these sequences suggests that predicting TR rates requires a detailed understanding of the nucleotide context and further experimental evidence. Additionally, our findings highlight the importance of both upstream and downstream sequences in modulating TR efficiency, with potential implications for therapeutic strategies targeting TR.

## Materials and Methods

### DNA constructs

Oligonucleotides containing sequences complementary to the corresponding SCC, with 10 nucleotides upstream and downstream of the stop codon, and overhangs matching the BspEI and BstEII restriction sites were designed. Dual reporter plasmids contained different SCCs and were constructed by annealing (1 µM each) oligonucleotides (Sigma) OST3230-3249 and OST3506-3541 (Supplementary Table 1, Supplementary Table 2) in an annealing buffer (1 mM EDTA in diethylpyrocarbonate-treated H_2_O, 100 mM NaCl, 10 mM Tris/HCl pH 7.5) in a thermocycler. Oligonucleotides were denatured at 98°C for 5 seconds, annealed at 40°C for 5 seconds, and then kept at 10°C. Annealed oligonucleotides were inserted into BspEI and BstEII sites of pcDNA3.1(+)RFP_MCS_GFP (PST1596) [41].

### Cell Culture

HeLa cells were cultured in high glucose Dulbeccós Modified Eagle Medium essential medium (DMEM, Gibco) supplemented with 10% heat-inactivated fetal calve serum (v/v) (Biowest), 1% L-glutamine (w/v), 100 µg/mL penicillin/streptomycin (P/S) at 37°C, 5% CO_2_ and 90% humidity. 30,000 cells/well were seeded in 96-well plates for the dual reporter assay.

Cells were transfected with 150 ng of plasmid DNA using Effectene (Qiagen) following the manufacturer’s instructions. The transfection reagent was removed 16 hours after transfection. When indicated, geneticin (Carl Roth) was added at 100 µg/mL for 24 hours.

### Flow Cytometry

The cells were cultured in a 96-well plate and prepared flow cytometry as previously described [41]. Briefly, the medium was discarded, and transfected cells were washed with 150 µL PBS. Cells were then trypsinized with 35 µL 0.5% trypsin and incubated for 10 minutes at 37°C and resuspended in 165 µL DMEM. Cells were pelleted for 5 minutes at 500 x g (Megafuge 8 centrifuge, ThermoFisher Scientific). The medium was replaced by 200 µL phenol red-free DMEM before flow cytometry (Guava easyCyte, Cytec). Cells were gated by forward scatter between 20,000 and 75,000. Side scatter was set between 7,000 and 60,000. Additionally, the cell population was gated for fluorescence signals (488 nm and 561 nm lasers) based on generating a two-parameter dot-plot of tagRFP detection vs. eGFP detection. The following thresholds were applied: RFP > 700 and GFP > 100. Up to 10,000 gated events were recorded over a 180 s interval per sample. For each biological replicate, three technical replicates were measured for each condition. For each measurement, a 100% TR control (cells transfected with PST1596) was measured in three replicates. Data was saved as a flow cytometry standard file (.fcs) for further analysis. To evaluate TR efficiency, the ratio of eGFP to tagRFP fluorescence normalized to the 100% TR control (PST1596). All gating and calculation steps were performed in RStudio utilizing the R programming environment.

### Statistics

Statistical analysis was done with Prism 10 (Graph Pad) using the one-was ANOVA and applying Dunnett’s test for multiple comparison correction. Data were presented as means + s.e.m. (standard error of the mean).

## Acknowledgments

We thank the members of the Thoms lab for the discussions and Maximilian Heppel and Sinan Zimmermann for comments on the mansucript. This research was supported by a grant from the German-Israeli Foundation for Scientific Research and Development (GIF, grant number 1459), a grant from the Deutsche Forschungsgemeinschaft (DFG) (TH 1538/3-1), and the Horst and Eva-Luise Köhler Foundation.

## Supplements

**Supplementary Figure 1:**
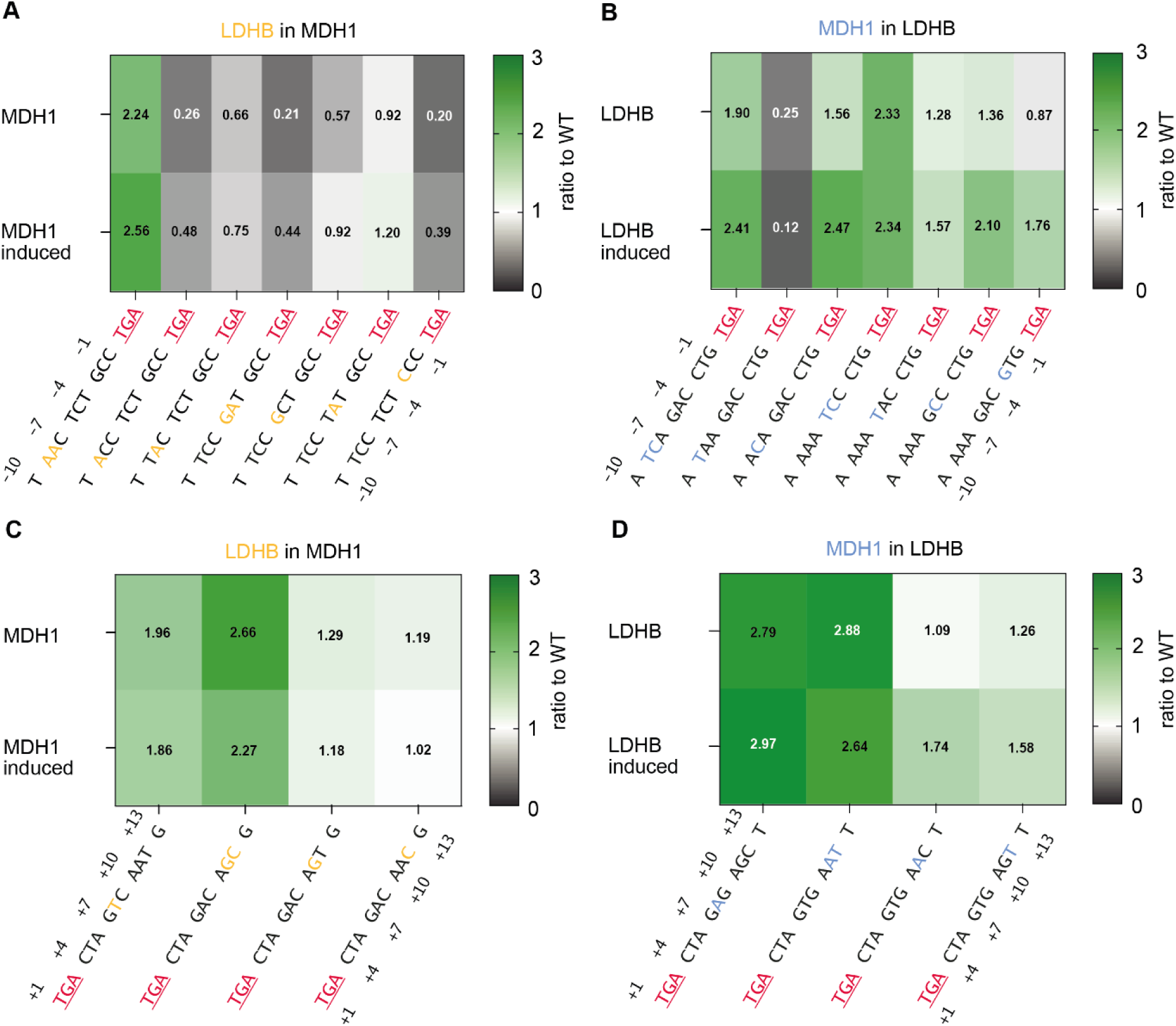
Ratios of Basal and G418-Induced TR: **A-D)** Heat map of TR ratios for WT SCCs of *MDH1* and *LDHB* and constructs with nucleotide exchanges. Green indicates increased TR, grey indicates decreased TR. Reference values set to 1. Calculations were done using PRISM. **A)** *MDH1* basal and induced TR for upstream sequences. **B)** *LDHB* basal and induced TR for upstream sequences. **C)** *MDH1* basal and induced TR for downstream sequences. **D)** *LDHB* basal and induced TR for downstream sequences. Measurements were conducted at least three times with three replicates.

**Supplementary Table 1:**
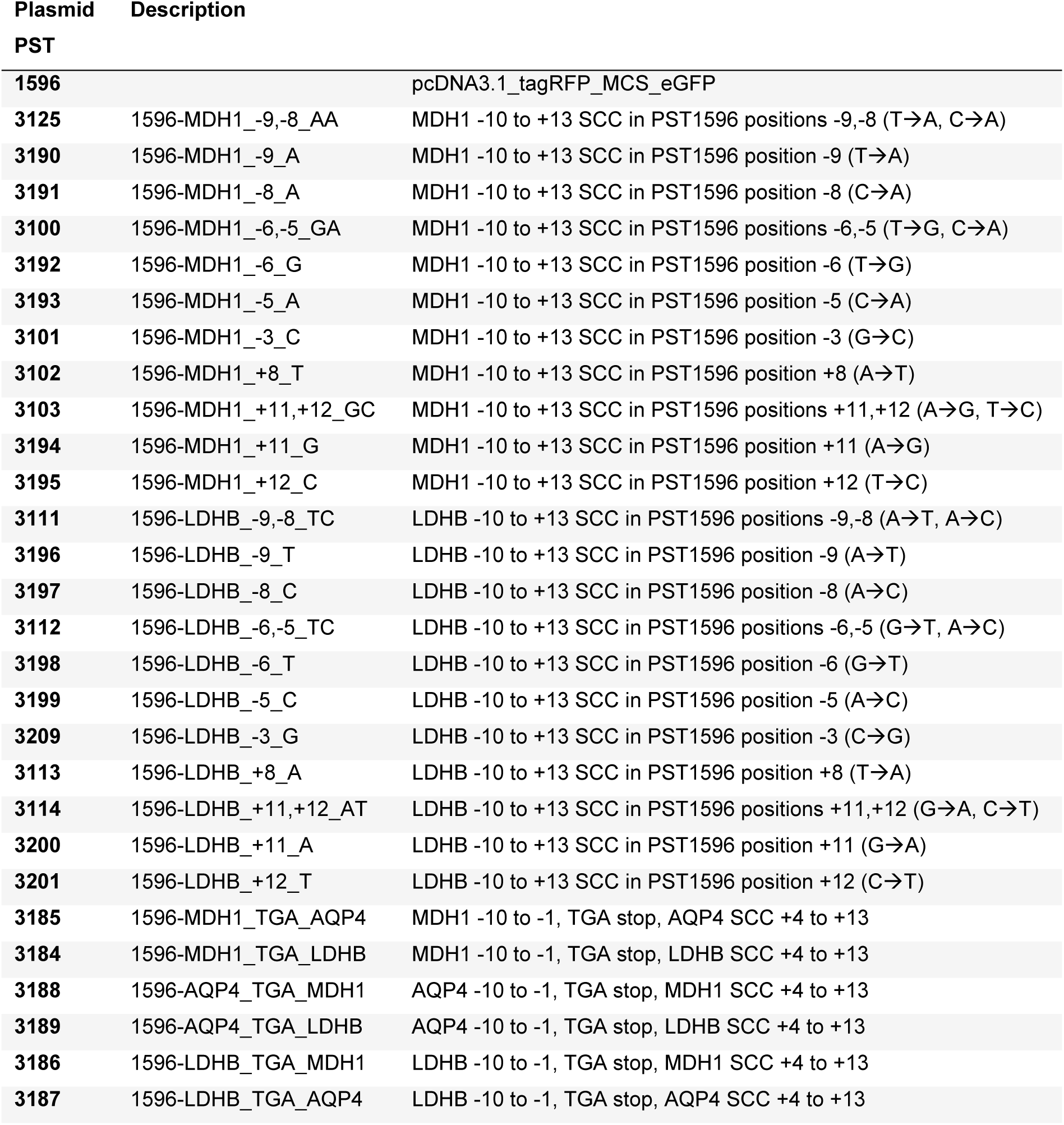
Plasmids used in this study.

**Supplementary Table 2:**
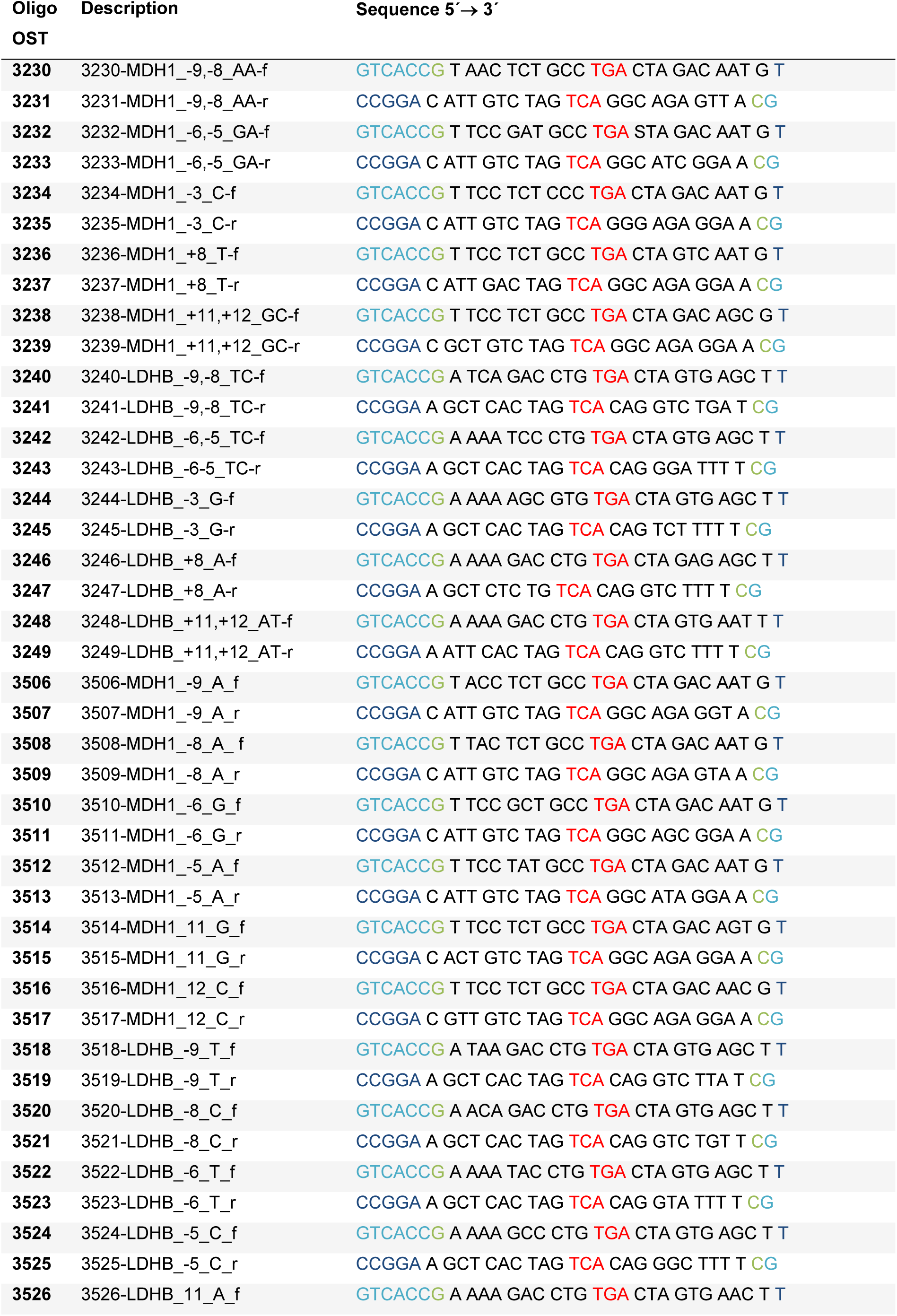
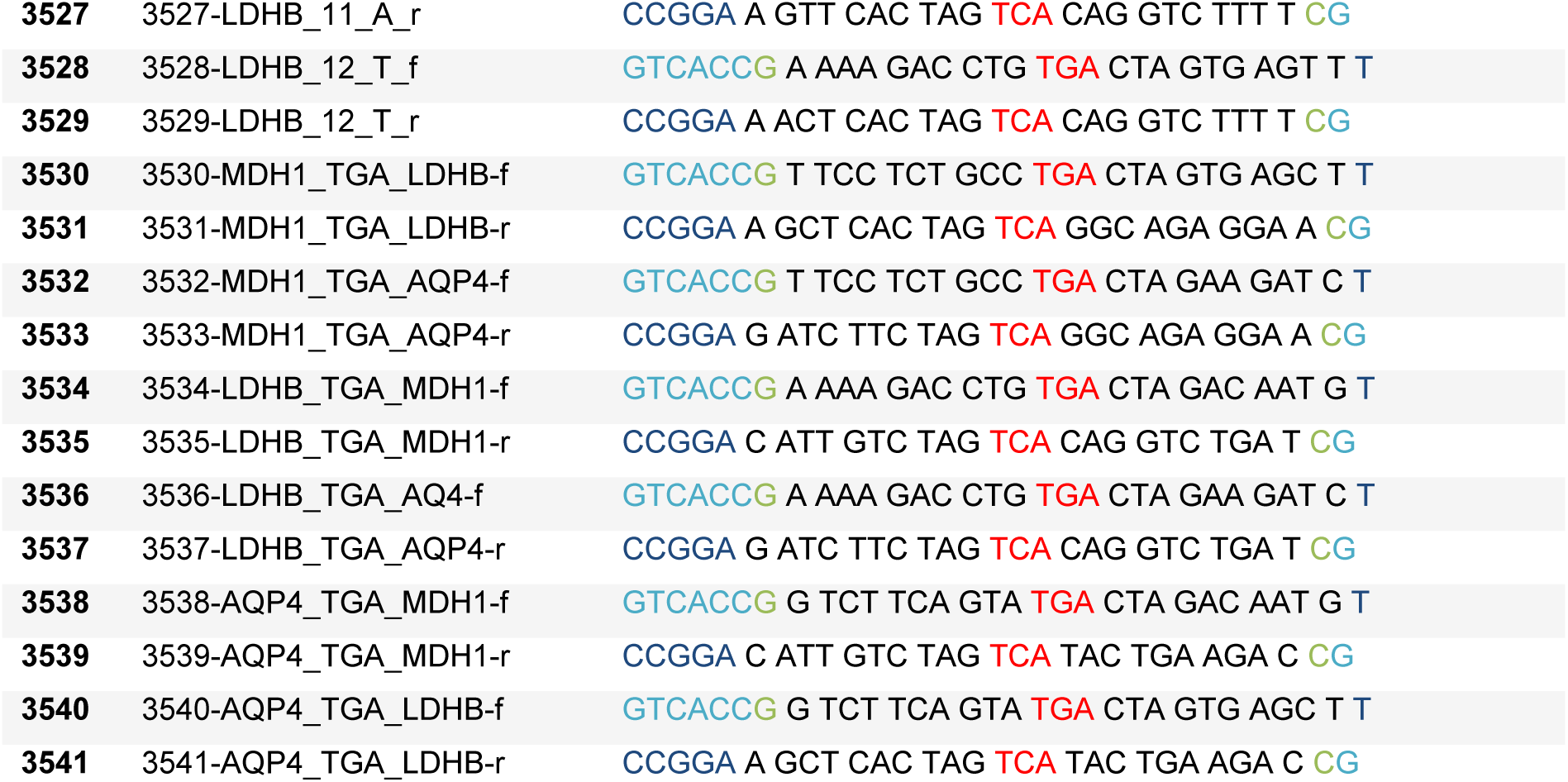
Oligonucleotides used in this study.

